# Pyruvate kinase M2 regulates Japanese encephalitis virus replication by interacting with NS1 protein

**DOI:** 10.1101/2024.03.12.584590

**Authors:** Vijay Singh Bohara, Atharva Deshmukh, Sachin Kumar

## Abstract

Pyruvate kinase isoform M2 (PKM2) is a key modulator of glucose metabolism. While the major role of PKM2 is to facilitate the breakdown of glucose, it is potentially associated with other additional non-glycolytic functions as well. The role of PKM2 in the autoimmune response and inflammatory process is increasingly being acknowledged as a crucial modulator of cellular pathophysiological activity. However, its role in modulating viral replication has not been explored in detail. In the present study, we have shown a significant increase in endogenous PKM2 expression in JEV-infected mouse neuroblastoma cells. Furthermore, overexpression of PKM2 significantly reduced JEV replication, suggesting a negative effect of PKM2 on JEV replication. This was further confirmed by siRNA-mediated downregulation of endogenous PKM2 expression, which resulted in enhanced JEV replication*. In silico* studies revealed the potential interaction between PKM2 and NS1 protein of JEV. The microscopic studies also showed cellular colocalization of PKM2 and NS1 in the ER of infected cells. The interaction was further validated *in vitro* by co-immunoprecipitation assay. The present study suggests that PKM2 negatively regulates the JEV replication by its possible interaction with NS1.

**Importance:** Japanese encephalitis (JE) is a neuroinflammatory disease caused by the Japanese encephalitis virus (JEV). JE is a major threat to public health not only because it causes many deaths but also for its permanent neuropsychiatric sequelae in children. Out of all non-structural proteins of JEV, NS1 is highly immunogenic. A wide range of possible interactive partners has been identified for the NS1, many of those have been linked to immune evasion and regulating viral replication. In the current study, we have described a novel host cell factor, PKM2 modulating JEV replication by interacting with NS1 protein. Considering PKM2’s central role in regulating host cell metabolism, our findings suggest a previously unrecognized role for PKM2 in JEV neuropathogenesis. The identification and characterization of previously unknown host factors, as well as the elucidation of their regulatory mechanisms, are of utmost importance in the development of innovative treatments and antivirals against JEV.

## Introduction

The Japanese encephalitis virus (JEV) is the causative agent of Japanese encephalitis, a prominent kind of viral encephalitis prevalent in Asia (1). JEV is classified as a member of the genus Flavivirus under the family flaviviridae (2). The positive sense 10.7 kb single-stranded RNA genome of the JEV, (2, 3) encodes a single polyprotein that is processed into three structural proteins (capsid, pre-membrane, and envelope) and seven non-structural (NS) proteins (NS1, NS2A/B, NS3, NS4A/B, and NS5) by viral and host proteases (4). The replication of JEV genomic RNA occurs in the viral “replication complex” that relates to membranes produced from the endoplasmic reticulum (ER) (5). The replication process is facilitated by the JEV non-structural proteins, in conjunction with various host factors (6–8). Host factors employ diverse strategies to both support or impede virus replication. It is therefore critical to elucidate the host factors that influence the replication cycle of JEV and to investigate the underlying molecular mechanism. The intricate role of NS1 in the host defense and viral replication makes it an attractive protein to study

Pyruvate kinase muscle isozyme M2 (PKM2) is a limiting glycolytic enzyme essential for tumor metabolism and development (9). PKM2 catalysis the conversion of phosphoenolpyruvate to pyruvate, the last ATP generating step of glycolysis (10). PKM2 also plays a pivotal role in multiple cellular pathways, encompassing aerobic glycolysis, intranuclear signal transmission, protein synthesis, inflammation, and apoptosis (11). PKM2 enhances the expression of genes by activating hypoxia-inducible factor-1α (HIF-1α), β-catenin (β-cat), insulin, signal transducers and activators of transcription 3 (STAT3), and other transcription factors that stimulate cell growth and proliferation (12–14). PKM2 also serves as a critical regulator in the apoptotic signaling pathways of several cancer types. B-cell lymphoma 2 (BCL-2), a prominent member of the BCL-2 family and very well characterized for its anti-apoptotic properties, is a target of PKM2, both directly and indirectly (15). As part of the innate immune response, PKM2 stimulates the production of inflammatory cytokines such as interleukin 1β (IL-1β) and tumor necrosis factor-α (TNF-α) (16). The PKM2 also functions as a protein kinase, facilitating the phosphorylation of the STAT3. This process then leads to an increase in the production of interleukin-6 (IL-6) and interleukin-1 beta (IL-1β), thereby initiating systemic tissue inflammation (17). PKM2 activates and engages with HIF-1α to regulate the release of high mobility group box-1 (HMGB1), an efficient proinflammatory cytokine that is released from activated macrophages (18, 19). Recent studies have shown an increase in PKM2 expression in several disorders that are characterized by the presence of autoimmune and inflammatory mediators (20–22). PKM2 performs several non-glycolytic tasks with far-reaching consequences, the extent of which has to be fully deciphered (23). Hence, it is important to have a more comprehensive understanding of the metabolic and nonmetabolic roles of PKM2.

PKM2 has been associated with several viral infections. The expression of PKM2 was enhanced in mouse lung tissues infected with H1N1 (24). PKM2 has been demonstrated to exhibit elevated expression levels in individuals affected with severe coronavirus disease 2019 (COVID-19) (25). Collectively, these studies suggest the potential involvement of PKM2 in viral pathogenesis. However, there has been a lack of research conducted specifically on flaviviruses, particularly the Japanese encephalitis virus (JEV).

The present study focuses on investigating the potential role of PKM2 in JEV neuropathogenesis. This was accomplished by examining the effects of overexpression of PKM2 and silencing of endogenous PKM2 on JEV replication in mouse neuroblastoma cells. *In-silico* and microscopic analyses were also performed to investigate the potential interaction between PKM2 and NS1. This was further confirmed in vitro using the co-immunoprecipitation (co-IP) technique.

## Materials and Methods

### Cells and Virus

The mouse neuroblastoma Neuro-2a, and the baby hamster kidney BHK-21, were obtained from the National Centre for Cell Sciences (NCCS), Pune, India. The cells were cultured in Dulbecco’s modified eagle medium (DMEM) with 10% fetal bovine serum (FBS) and 1% antibiotic-antimycotic (GIBCO) cocktail at 37 °C humidified incubator in 5% CO2. The JEV strain SA14-14-2 (GenBank accession number JN604986) used in this study was procured from the Department of Health and Family Welfare, Government of Assam, India. The JEV stock preparation and its titration by plaque assay were performed in BHK-21 cells. For stock preparation, Neuro2a cells at 80 % confluency, were infected with JEV at 0.1 MOI. Following a period of 2 hours during which the JEV was allowed to adsorb to the cells, the infection media was subsequently substituted with DMEM media supplemented with 2% fetal bovine serum (FBS). Cells were lysed in successive freeze-thaw cycles 72 hours post-infection. Cell debris was removed by centrifuging at 1,500 g for 5 minutes, clear supernatant was collected and stored at -80 °C. Later, the presence of JEV was confirmed by plaque assay.

### Plasmids and Antibodies

The following plasmids were used in the current study: pEGFP.PKM2 (Addgene #64698) and pEGFP (Addgene #165830). The antibodies used are as follows: Anti-NS1 (GeneTex, USA), Anti-PKM2 (Cell Signaling Technology, USA), Anti-GAPDH (Cell Signaling Technology, USA), Anti-GFP (Bio Bharati, India), Goat anti-Rabbit (Invitrogen, USA) and anti-mouse (Cell Signaling Technology, USA) conjugated with horseradish peroxidase (HRP).

### Virus Infection and Quantification

Neuro2a cells were seeded in a 6-well plate at a density of 0.6 X 10^6^ cells/ well and infected with JEV at 0.1 MOI. The virus was allowed to adsorb for 2 hours with intermittent shaking. Thereafter, infection media was replaced with fresh DMEM (2 ml/well) supplemented with 2% FBS and the plate was placed in a 5% CO_2_ incubator. Cells and Supernatants from JEV-infected cells were collected at different time points post-infection. Collected cells were used for expression analysis of endogenous PKM2 and JEV NS1 protein using quantitative real-time PCR and immunoblotting. The collected supernatant was used for virus quantification using Plaque assay as per standard protocol.

### Gene expression analysis using Real-time PCR

RNA lysate was prepared at different time points post-infection using RNAiso Plus reagent (TaKaRa, Japan). Total RNA was extracted using the phenol-chloroform extraction method and 1 µg of total RNA was reverse transcribed into cDNA using a high-capacity cDNA reverse transcription kit (Thermo Fisher Scientific, USA). The quantitative real-time PCR was performed using PowerUp SYBER Green Master mix (Applied Biosystems, USA). The fold change in mRNA level in infected versus mock-infected samples was calculated using the 2^(-ΔΔCt)^ method keeping GAPDH as an internal control for normalization.

### Immunoblotting

Cells were lysed in RIPA buffer. An equal volume of proteins was loaded and separated on 12% SDS gel. Separated proteins from the gel were transferred to a 0.4 µm nitrocellulose membrane. The membrane was blocked with 5% skimmed milk prepared in 1X TBS buffer in 0.1% TBST for 2 hours. Following blocking, primary antibody dilution was prepared in 2% BSA (Bovine Serum Albumin) and incubated at 4°C overnight. After incubation, the membrane was washed with 1X PBS and thereafter incubated with HRP-conjugated secondary antibody for 1 hour at room temperature and detected using ECL reagent (BioRad, USA) under chemiluminescence. Quantitative analysis was performed using ImageJ software.

### Overexpression and Knockdown studies

The plasmid used in this study pEGFP-PKM2 (Addgene #64698) is a generous gift from Axel Ullrich lab. For overexpression experiments, Neuro-2a cells were cultured on 6-well plates and then transfected with a total of 2 µg of an expression plasmid using Lipofectamine 2000 (Invitrogen, USA) as per the instructions provided by the manufacturer.

The knockdown of endogenous PKM2 was achieved by co-transfecting two siRNAs at a total concentration of X pmol. The following sense strand sequences of siRNA were used: GAUGUCGACCUUCGUGUAA[dT] and UCCUAUCAUUGCCGUGACU[dT][dT] (26). Lipofectamine RNAiMAX (Invitrogen, USA) was used for transfection studies as per the manufacturer protocol, and siRNA universal negative control #1 (SIC001, Sigma-Aldrich, Germany) served as a negative control.

### Structure modeling of JEV NS1 and mouse PKM2 protein

The amino acid sequences of JEV NS1 and mouse PKM2 were retrieved from GenBank accession numbers, QCZ42158 and NP_001365797, respectively. With the help of these sequences, three-dimensional structures of the proteins were predicted and generated by I-TASSER (Iterative Threading ASSEmbly Refinement) protein structure & function prediction software. Further, this model was validated by PROCHECK-Ramachandran plot after its energy minimization using Yasara software.

### Molecular docking study

The interaction between the predicted structure for JEV NS1 and mouse PKM2 proteins was performed by molecular docking. Docking was performed using Cluspro 2.0 protein-protein docking software. The PDB sum generator server was used to find the interacting amino acid residues spanning both proteins. Finally, the top docked models obtained by Cluspro 2.0 were analyzed and ranked based on the MM/GBSA free energy decomposition with the help of the HawkDock server.

### Molecular dynamics simulations

The docked protein-protein complex was subjected to MD simulations to study the conformational stability using the GROMACS v2020.1 software program to obtain the parameters of proteins. The system consisted of the protein-protein complex in a solvated dodecahedron box with a minimum distance of 1.2 nm from the boundary. The system was filled with single-point charge water and subsequently neutralized by adding countercations (Na^+^) or anions (Cl^-^). The solvated system was then energy minimized using the steepest descent method, followed by the equilibrium for 100 ps through NVT and NPT ensembles to optimize the orientation and system density. The final equilibrated system was used as starting conformations to run the MD simulations for 50 ns. Finally, the output trajectory was obtained, and the estimation of Root Mean Square Deviation (RMSD), Root Mean Square Fluctuation (RMSF), Solvent Accessible Surface Area (SASA), and Radius of Gyration (Rg) was done using GROMACS packages. The graphs were analyzed and plotted using XMGRACE software.

### Immunofluorescence

Neuro-2a cells were grown on coverslips and subsequently transfected with pEGFP-PKM2. The cells were infected with JEV 24 hours post-transfection and 48 hours post-infection, cells were fixed with 4% formaldehyde in 1X PBS for 15 minutes at RT. Following fixation, cells were washed with 1X PBS at least 3 times and permeabilized with 0.3% Triton X-100 in 1X PBS. Subsequently, the fixed cells were rinsed using 1X PBS and then incubated in a blocking buffer (5% BSA). For JEV protein detection, cells were incubated with 1/100 dilution of mouse monoclonal antibody against NS1 (GeneTex, USA) followed by incubation with 1/500 dilution of goat Alexa-fluor 594 (Invitrogen, USA) anti-mouse IgG secondary antibody. After the final wash with PBS, cells were stained with 4,6-diamidino-2-phenylindole dihydrochloride (DAPI). After final washing, slides were mounted in glass slides using prolong Gold Antifade Reagent (Invitrogen, USA) and examined under a 63X objective using the confocal microscope.

### Co-immunoprecipitation (Co-IP) assay

Capturem Co-IP kit was procured from TaKaRa (Japan). The Neuro-2a cells were transfected with pEGFP-PKM2 and cells were infected with JEV 24 hours post-transfection. Infected cells were harvested and lysed after 48 hours in the 200 µl lysis buffer per 1 x 10^6^ cells. An appropriate amount of the protease inhibitor cocktail was added to the lysis buffer to yield a 1X final concentration. The collected lysate was centrifuged at 17,000g for 10 minutes at 4°C. The clear supernatant was collected and incubated with the recommended amount of anti-GFP (Bio Bharati, India) and anti-NS1 (GeneTex, USA) antibodies for 3 hours at 4°C. Pre-incubated samples were loaded onto the Capturem protein A columns (TaKaRa) and centrifuged at 1000g for 1 minute at room temperature. Flowthrough was discarded and spin columns were washed with wash buffer (provided in the kit). Bound samples were eluted from spin columns in elution buffer by centrifugation at 1000g for 1 minute at room temperature and analyzed by western blot.

### Statistical Analysis

All the data were statistically validated through two-tailed analyses using Student’s t-test using GraphPad Prism software, and results were shown as mean ± standard deviation. The level of significance among different groups was presented as *, **, *** where * is for P < 0.05, ** for P < 0.01, and *** for P < 0.001. P value less than 0.05 was considered significant.

### Data availability

The data sets generated in the study are available from upon appropriate request from the corresponding author.

## Results

### JEV infection induces upregulation of PKM2 expression in Neuro-2a cells

The expression of PKM2 was investigated in Neuro-2a cells following JEV infection with different multiplicities of infection (MOI). Quantitative analysis using real-time PCR at various MOI showed a gradual increase in JEV NS1 mRNA (Figure 1A). PKM2 mRNA also showed an increase in expression compared to control with the increase in MOI except at 0.001 MOI (Figure 1B). However, the maximum increase in expression was at 0.1 MOI. At the protein level, PKM2 expression was significantly enhanced with an increase in MOI (Figure 1C). Further, the time-dependent experiment was performed to analyze PKM2 expression post-JEV infection. The time-dependent fold-change in JEV NS1 mRNA was also examined (Figure 1D). The mRNA analysis revealed an upregulation of PKM2 expression at both 24- and 48 hours post-infection. There was no significant difference in the PKM2 mRNA compared to control at 72 hours post-infection (Figure 1E). At the protein level, there was a significant increase in PKM2 expression in infected cells compared to control (Figure 1F).

**Figure 1.**
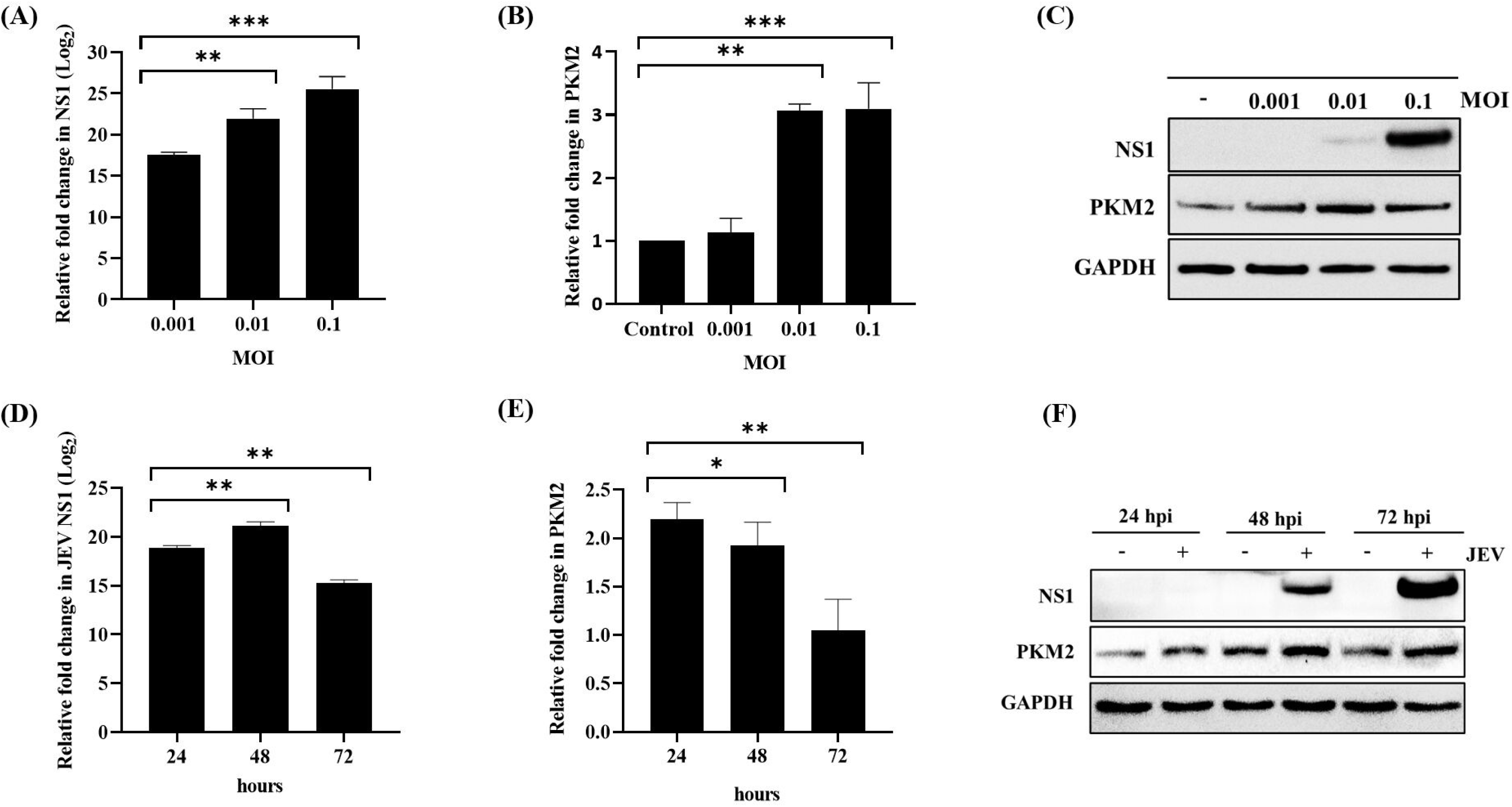
JEV infection leads to upregulation of PKM2. Real-time PCR analysis of JEV infection at different MOI post-infection (A). Fold-change in PKM2 mRNA at different MOI post-infection (B). Immunoblot showing MOI-dependent expression of PKM2 and JEV infection at protein level (C). Time-dependent analysis of JEV infection at mRNA level (D). Time-dependent fold change in PKM2 mRNA expression post-infection (E). Immunoblot analysis of PKM2 expression and JEV infection at different time points post-infection (F). For Real-time PCR analysis, GAPDH was used for normalization.

### PKM2 expression is responsible for inhibiting JEV replication

To investigate the function of PKM2 in the context of JEV infection, neuro-2a cells were transfected with pEGFP-PKM2 plasmid (Figure 2A). PKM2 expression was confirmed at both 24- and 48 hours post-transfection (Figure 2B). Next, the cells were infected with JEV at 0.1 MOI, 24 hours post-transfection, and protein lysate was prepared 48 hours post-infection. The western blot analysis revealed a 0.394-fold reduction in JEV infection in cells with over-expressed PKM2 compared to untransfected cells (Figure 2C). This was further validated in an immunofluorescence experiment that showed a similar reduction in JEV infection in cells with overexpressed PKM2 (Figure 2D). Using a standard plaque assay, the virus titer in the supernatant collected from the infected cells was also quantified (Figure 2E). It was observed that the JEV titer was reduced by 0.46-fold in cells overexpressing PKM2 compared to the control (Figure 2F).

**Figure 2.**
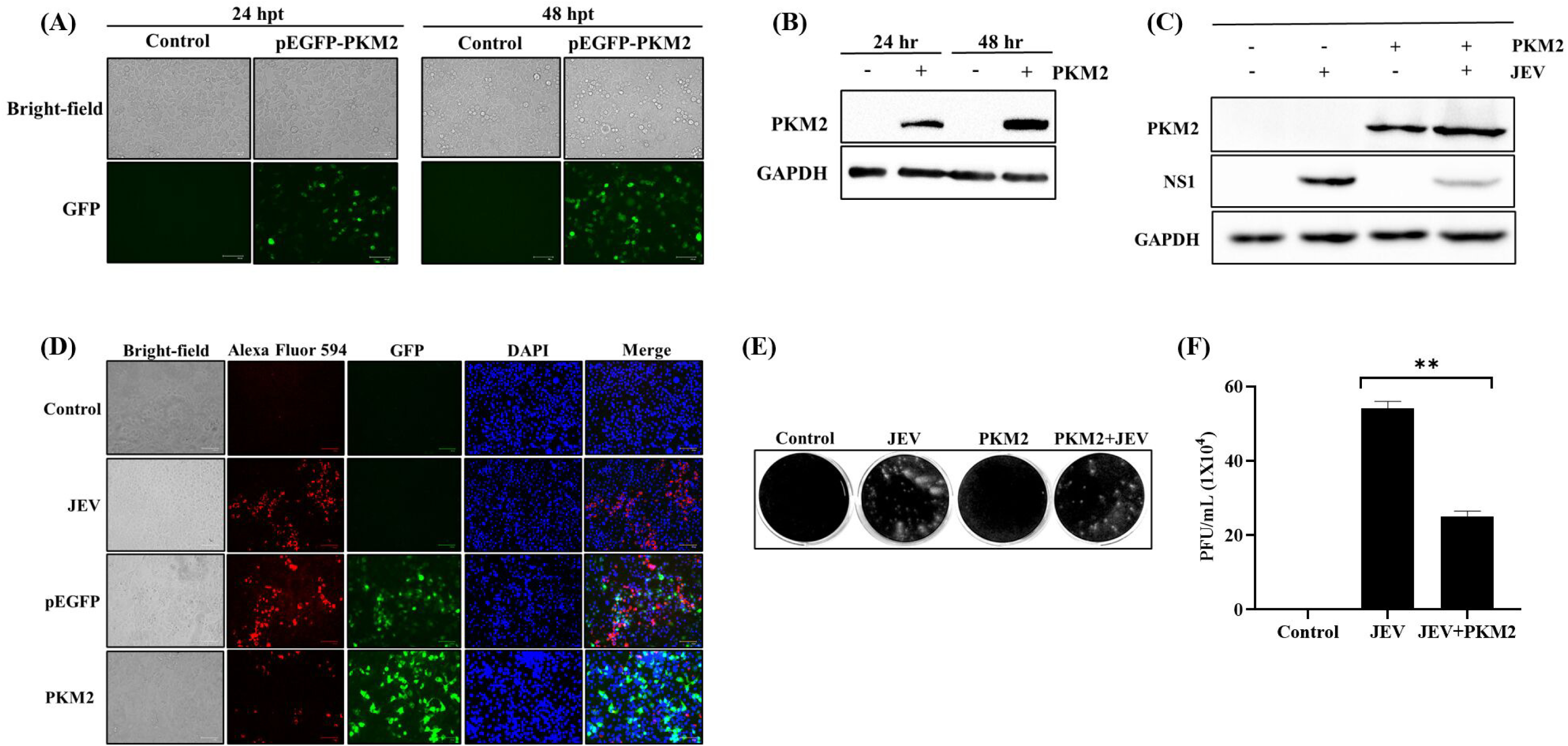
PKM2 overexpression inhibited JEV replication. Image shows expression of exogenous PKM2 24- and 48-hours post-transfection (A). Immunoblot analysis of time-dependent expression of exogenous PKM2 post-transfection (B). Immunoblot analysis of JEV infection upon overexpression of PKM2 (C). Immunofluorescence pictures of cells with exogenous PKM2 expression and JEV infection. Immunofluorescence labeling of JEV; red fluorescence indicates JEV-infected cells; blue fluorescence indicates DAPI-labelled nuclei; and green fluorescence represents PKM2 expression (D). Picture showing the reduction in extracellular viral titer in the form of plaques (E). Graph representing virus titer level measured in plaque-forming units per millilitre (PFU/ml) (F).

### PKM2 downregulation enhanced JEV replication

Endogenous PKM2 was silenced via transfection of a cocktail consisting of two PKM2-specific siRNAs. To quantify the PKM2 downregulation, real-time PCR and western blot analyses were performed. At the mRNA level, a 0.16-fold reduction in PKM2 expression was observed as compared to -ve siRNA-transfected cells (Figure 3A). At the protein level, the transfection of specific siRNA resulted in a 0.002-fold reduction in PKM2 expression compared to -ve siRNA transfected cells (Figure 3B). The western blot analysis demonstrated around a 1.8-fold increase in JEV infection in cells exhibiting downregulated PKM2, as compared to -ve siRNA transfected cells (Figure 3C). Immunofluorescence results also showed enhanced JEV infection in PKM2-silenced cells (Figure 3D). Extracellular virus was titrated using plaque assay (Figure 3E). JEV titer in supernatant collected from PKM2 downregulated cells was 3.6x10^5^ pfu/ml. In the supernatant of untransfected and -ve siRNA transfected cells, titer was 1.7x10^5^ pfu/ml and 1.4x10^5^ respectively. When compared to -ve siRNA transfected cells, supernatant from PKM2 silenced cells showed 2.56-fold higher JEV titer and compared to untransfected cells titer was 2.1-fold higher (Figure 3F).

**Figure 3.**
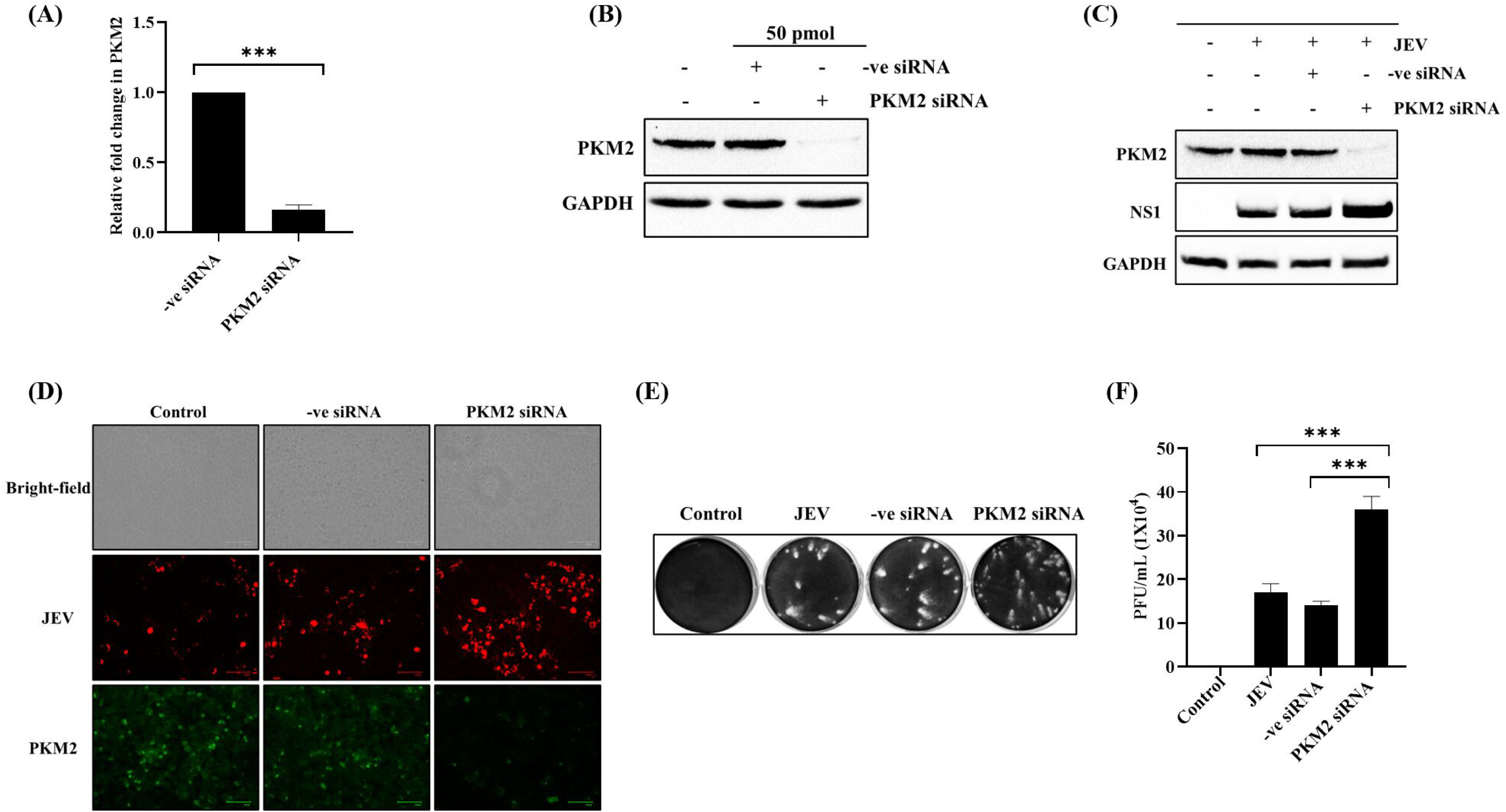
PKM2 knockdown enhanced JEV replication. Immunoblot showing the reduction in PKM2 mRNA level upon treatment with PKM2 siRNA (A). Immunoblot analysis of JEV infection upon knockdown of PKM2 (B). Graphs showing extracellular viral titer in the form of plaques in cells treated with PKM2 siRNA compared to untreated and -ve siRNA-treated cells (C). Immunofluorescence images showing enhanced JEV infection upon treatment with PKM2 si-RNA. Red fluorescence images indicate JEV-infected cells while green fluorescence indicates endogenous PKM2 expression (D). Plaque images show the reduction in extracellular viral titer (E). Graph representing virus titer level in plaque forming units per millilitre (PFU/ml) (F).

### Structure modeling of JEV NS1 and mouse PKM2 protein

The three-dimensional structure of mouse PKM2 protein was modeled (Figure 4A). The modeled structure was analyzed using a Ramachandran plot which showed 90.8% residues in most favoured regions, 7.5% in additional allowed regions, 0.9% in generously allowed regions, and only 0.9% in disallowed regions (Figure 4B). Similarly, the three-dimensional model of the JEV NS1 protein was predicted (Figure 4C). The model was analyzed using a Ramachandran plot and showed 87.8% residues in most favoured regions, 11.6% in additional allowed regions, 0.3% in generously allowed regions, and only 0.3% in disallowed regions (Figure 4D).

**Figure 4.**
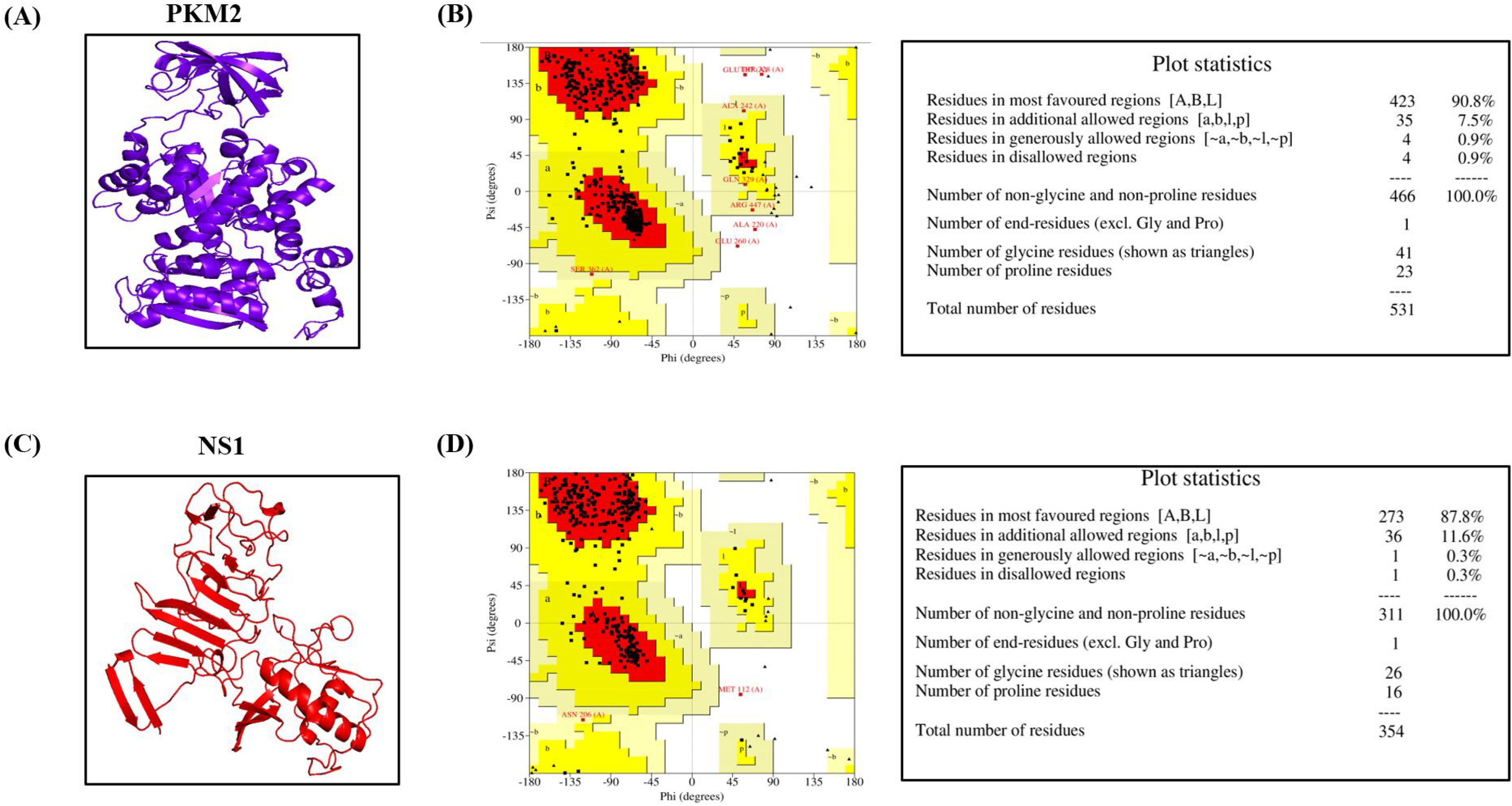
Structure modeling of JEV NS1 and mouse PKM2 protein. Figures depicting the three-dimensional model of Mouse PKM2 protein (A) with its analyzed Ramachandran plot (B) and the three-dimensional structure of JEV NS1 protein (C) with its analyzed Ramachandran plot (D).

### PKM2-NS1 Docking studies

Docking between the mouse PKM2 protein and NS1 protein of JEV was performed (Figure 5A). The integrity of the docked complex model was analyzed using PROCHECK. The Ramachandran plot of the PKM2-NS1 protein complex showed 647 residues (83.3%) in the most favourable regions, 112 residues (14.4%) in additionally allowed regions, 10 residues (1.3%) in the generously allowed regions, and 8 residues (1.0%) in the disallowed regions (Figure 5B). The interface residues of the complex were also analyzed. The 33 residues of PKM2 interacted with 33 residues of NS1 (Figure 5C). The number of salt bridges was 6, hydrogen bonds were 24, and non-bonded contacts were 372 (Table 1).

**Figure 5.**
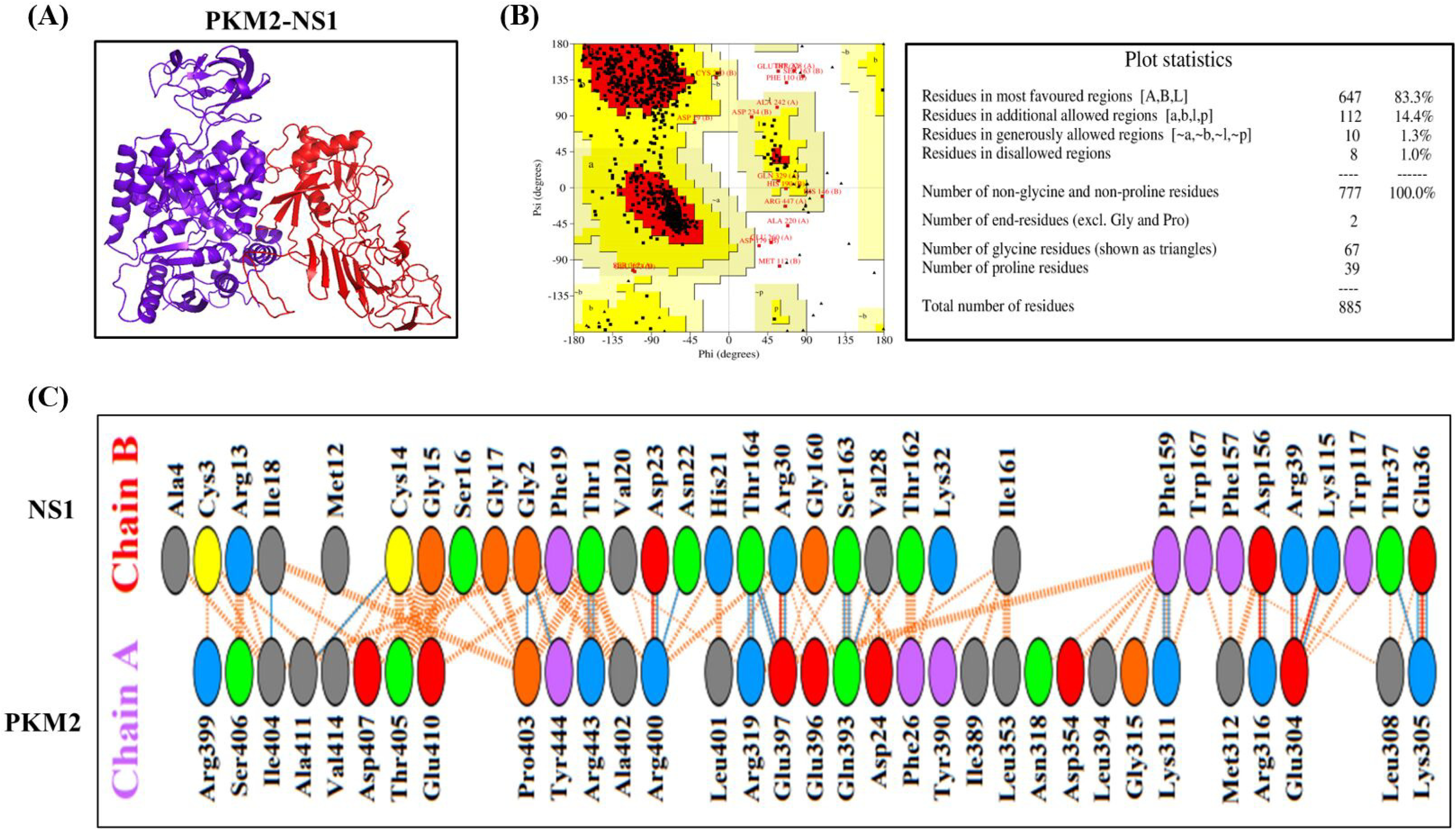
PKM2-NS1 Docking studies. Visualization of the PKM2-NS1 docked complex (A) with its analyzed Ramachandran plot (B) and their interacting amino acid residues (C).

**Table 1.**
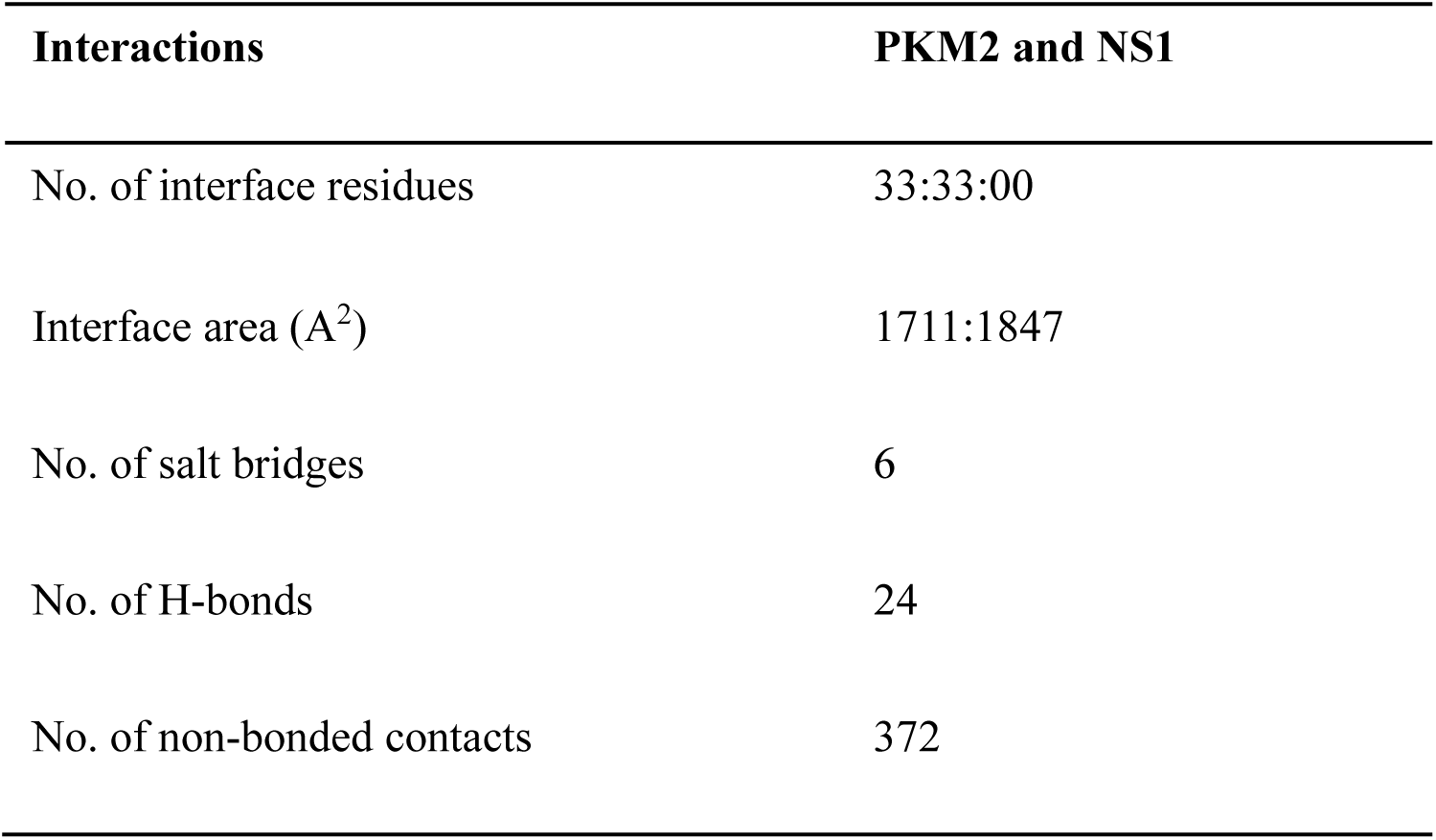
The number of interface residues, interface area, salt bridges, H-bonds, and non-bonded contacts that exist between PKM2 and NS1 interaction.

| **Interactions** | **PKM2 and NS1** |
| --- | --- |
| No. of interface residues | 33:33:00 |
| Interface area (A<sup>2</sup>) | 1711:1847 |
| No. of salt bridges | 6 |
| No. of H-bonds | 24 |
| No. of non-bonded contacts | 372 |

### Molecular dynamics simulations

MD simulations of JEV NS1 complexed mouse PKM2 protein for a 50 ns timeframe were conducted to evaluate the binding strength and stability of the complex. Several parameters, including RMSD, RMSF, SASA, and Rg, were performed. The RMSD plot analysis is crucial in understanding the structural stability of protein and protein-ligand bound complex. The PKM2 and its complex achieved steadiness at 12 ns and 17 ns, and both kept its equilibrium with only minor changes. The average RMSD values obtained for PKM2 and its complex with NS1 were 0.35 nm and 0.325 nm respectively (Figure 6A). RMSF of the complex was also analyzed to evaluate the rigidity and flexibility of bound complexes. C-alpha atom of the residues of the bound complexes were analyzed to identify the fluctuations of each atom across the backbone. The PKM2 and its complex with NS1 exhibited stable fluctuations throughout, with minor fluctuations around 500 residue region (Figure 6B). The average RMSF value obtained for the PKM2 and its complex with NS1 were 0.119 nm and 0.154 nm respectively. Rigidity and compactness of the complex is characterized using Rg values. Rg values for PKM2 achieved equilibrium at 10 ns and maintained it till the end except for minor fluctuation around 20 ns and 35 ns, while PKM2-NS1 complex showed minor fluctuations at the start however achieved equilibrium at around 14 ns and maintained it till the end except for minor fluctuations at around 33–38 ns timeframe (Figure 6C). The average Rg value obtained for the PKM2 protein and its complex were 2.58 nm and 3.20 nm respectively. Additionally, the SASA value was analyzed to assess the solvent behaviour and examine the folding/unfolding of the PKM2 protein and its complex. The average SASA value obtained for the PKM2 protein and its complex with NS1 were 248.249 nm^2^ and 403.759 nm² respectively (Figure 6D).

**Figure 6.**
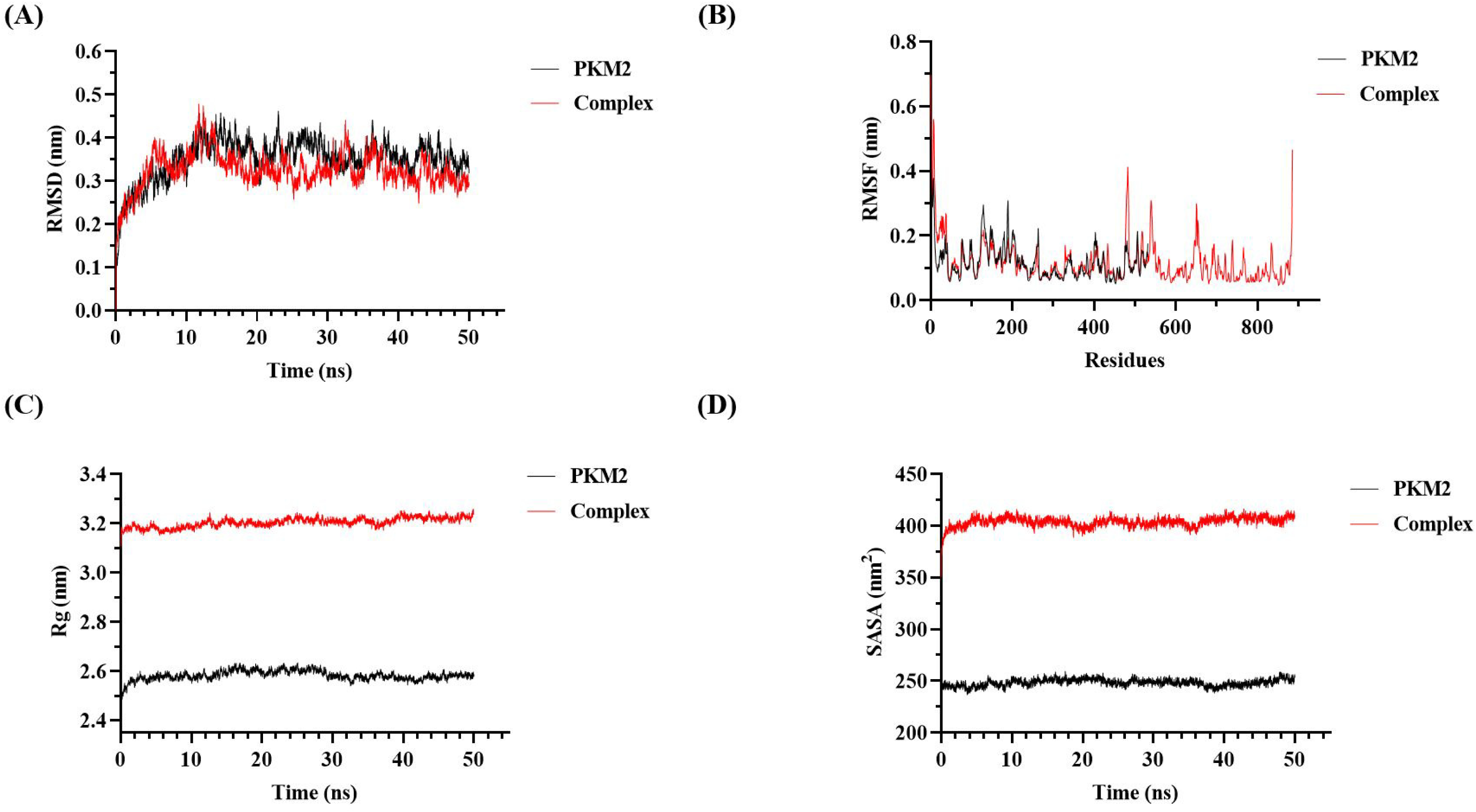
Molecular dynamics simulations. Graphs showing results after Root Mean Square Deviation (RMSD) analysis (A), Root Mean Square Fluctuations (RMSF) analysis (C), Radius of Gyration (Rg) analysis (B), and Solvent Accessible Surface Area (SASA) analysis (D).

### Cellular localization and interaction studies

Confocal microscopy was performed to study the cellular localization of PKM2 and NS1. Cells were transfected with PKM2 expressing plasmid and infected with JEV. Immunofluorescence results showed the cellular co-localization of PKM2 with NS1 protein in the ER of infected cells (Figures 7A and 7B). The Co-IP experiment was performed to study the interaction between PKM2 and NS1. PKM2 was detected after the pull-down with NS1, while NS1 was detected after the pull-down with GFP. This validated the interaction between PKM2 and NS1 protein (Figure 7C).

**Figure 7.**
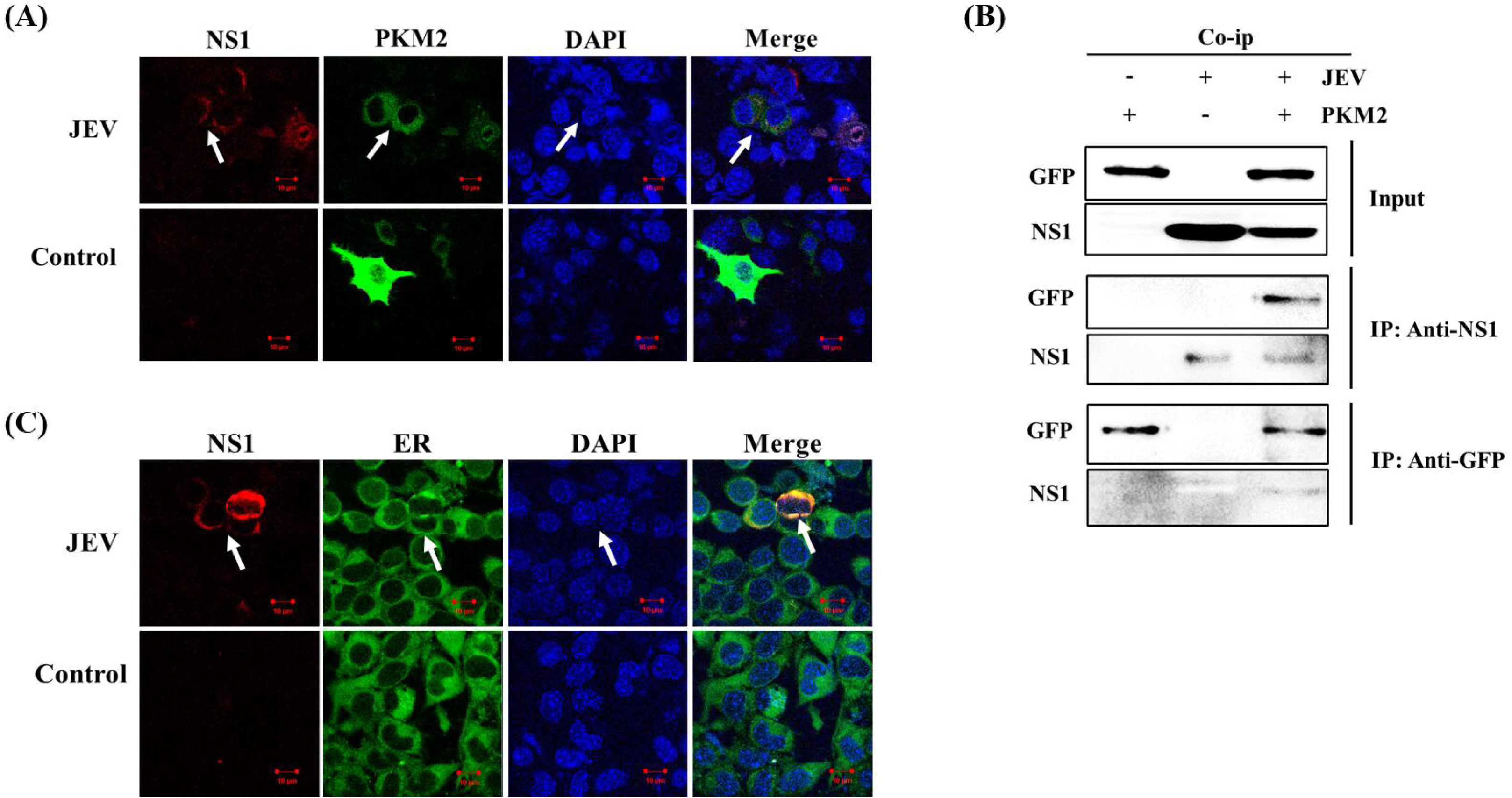
Cellular localization and interaction studies. Confocal images showing cellular co-localization of PKM2 with NS1 in JEV-infected cells. Representative images showing red fluorescence indicate NS1 expression while green fluorescence indicates PKM2 expression. The merge panel shows the co-localization of PKM2 and NS1 proteins. Cell nuclei were stained with DAPI (blue channel) (A). Confocal images showing localization of NS1 (red channel) and ER (green channel). ER was stained with ER tracker dye (Invitrogen) while the nucleus was stained with DAPI (B). The interaction of PKM2 with the NS1 protein of NDV was examined by co-immunoprecipitation assay (co-IP). The cell lysates were immunoprecipitated by anti-NS1 and anti-GFP antibodies and interactions were detected using immunoblot (C).

## Discussion

PKM2 is a glycolytic enzyme catalyzing rate limiting the last step of glycolysis (9). PKM2 is a multifunctional protein involved in several critical functions like apoptosis, mitosis, hypoxia, inflammation, and metabolic reprogramming (11, 27). The role of PKM2 in several cancers has been well explored (28–31). Furthermore, PKM2 participation in numerous cellular pathways, protein-protein interactions, and nuclear transport indicates that it performs several non-glycolytic functions (9). Perhaps, PKM2 might be involved directly or indirectly in regulating virus replication and pathogenesis through multiple pathways. Since the role of PKM2 in JEV neuropathogenesis has not been studied. The present work has explored a mechanism by which PKM2 might regulate JEV replication.

Firstly, MOI-dependent expression analysis of PKM2 was performed which showed an increase in its expression with the increase in MOI. Next, we performed a time-dependent analysis of PKM2 expression in JEV-infected Neuro-2a cells which showed higher PKM2 expression in infected cells compared to control cells at all time points post-infection. Both these studies hint towards the possible role of PKM2 in regulating JEV infection. To study the function of PKM2, we performed overexpression and knock-down studies. In the overexpression experiment, we transfected the cells with PKM2-expressing plasmid and then infected them with JEV. It was observed that JEV infection was significantly reduced in cells with overexpressed PKM2. We also examined the extracellular viral titer in the supernatant of infected cells and observed a significant reduction in JEV titer. Knockdown studies were also performed to further validate these results. We used PKM2-specific siRNA to silence endogenous PKM2 expression. The data suggests that JEV infection was enhanced in cells with silenced PKM2. Extracellular viral titer also showed enhanced virus particles in the supernatant collected from cells with silenced PKM2 compared to cells treated with -ve siRNA. All these results showed that PKM2 negatively affects JEV replication.

PKM2 is a sticky protein and is known to interact with several host and viral proteins. It has been observed that PKM2 exhibits interactions with the pathogenic E7 protein of HPV (human papillomavirus and NS5B of the hepatitis C virus (32–34). The viral RNA polymerase is another target of PKM2. These interactions of PKM2 with viral proteins positively or negatively modulate virus infection. NS1 protein is a non-structural protein that plays an important role in viral infection and propagation by contributing to viral replication and virulence, immunological invasion, and host complement system activation (35). Multiple studies have shown that NS1 is a necessary cofactor in the flavivirus RNA replication (36–39). It has been discovered that NS1 in cells is found near vesicle packets and cytoplasmic vacuoles, where dengue virus replication takes place in Vero and C6/36 cells (36). NS1 may play a structural role, together with other transmembrane replicase components, in anchoring the replication complex to the membrane (40–44). Experiments involving trans-complementation and mutagenesis in yellow fever virus or West Nile virus (WNV) have demonstrated that whatever function NS1 plays, it occurs early in RNA replication (37, 45). To elucidate the mechanism by which PKM2 regulates JEV replication molecular docking and simulations studies were performed.

The modeled mouse PKM2 and JEV NS1 protein structures using I-TASSER were used for docking studies using Cluspro 2.0. The docking studies revealed the possible protein-protein interaction. The analysis of the residues involved in the interaction showed the binding of NS1 protein with PKM2 together with the involvement of several salt bridges and hydrogen bond linkages. We also analyzed the rigidity and stability of the PKM2-NS1 complex with the help of MD simulations. The RMSD and RMSF patterns did not show any substantial shifts, which suggested the stability of PKM2 and its complex with NS1. Further, the increase in Rg and SASA values after NS1 binding can be explained due to increased protein-protein complex size. Also, the Rg and SASA patterns were overall consistent and suggested stable docked complex. Microscopic studies also showed cellular co-localization of PKM2 and NS1 protein suggesting their probable interaction. Further, the interaction was also validated by the Co-IP assay. However, other viral proteins might also interact with PKM2. Based on our results, we have inferred that one of the reasons for the suppression of JEV replication might be due to PKM2 and NS1 interaction.

JEV is an important neurotropic virus and displays a variety of neuropathophysiological alterations in the infected host. Although this study highlights the involvement of PKM2 in JEV neuropathology, there might be other host factors that could modulate its infectivity. This study paved a path to explore the possibility of using other host metabolic genes to elucidate the pathology of JEV in neurons.

## Conclusion

From the present study, we suggest PKM2 as a host factor that can modulate JEV replication by directly binding to the NS1 protein of JEV. Since PKM2 is gaining recognition as an essential regulator of cellular pathophysiological activity. Therefore, there could be other mechanisms through which PKM2 might be involved in JEV infection. Further studies will help in understanding the cross-talk between PKM2 and JEV infection.

## Acknowledgments

The funding for this work was provided by Department of Health Research, Government of India (Grant No.NER/71/2020-ECD-I).

## Author contributions

**Vijay Singh Bohara:** Conceptualization, Data curation, Formal analysis, Investigation, Methodology, Visualization, Writing - original draft. **Atharva Deshmukh**: Data curation, Formal analysis, Writing (Docking and Molecular dynamics simulations). **Sachin Kumar**: Conceptualization, Funding acquisition, Investigation, Project administration, Resources, Supervision, Validation, Writing- review and editing.

## REFERENCES

1. Solomon T. 2006. Control of Japanese encephalitis--within our grasp? N Engl J Med 355:869–71.

2. Turtle L, Solomon T. 2018. Japanese encephalitis — the prospects for new treatments. Nature Reviews Neurology 14:298–313.

3. Yun S-I, Lee Y-M. 2014. Japanese encephalitis. Human Vaccines & Immunotherapeutics 10:263–279.

4. Yun SI, Lee YM. 2018. Early Events in Japanese Encephalitis Virus Infection: Viral Entry. Pathogens 7.

5. Gillespie Leah K, Hoenen A, Morgan G, Mackenzie Jason M. 2010. The Endoplasmic Reticulum Provides the Membrane Platform for Biogenesis of the Flavivirus Replication Complex. Journal of Virology 84:10438–10447.

6. Paranjape SM, Harris E. 2010. Control of dengue virus translation and replication. Curr Top Microbiol Immunol 338:15–34.

7. Villordo SM, Gamarnik AV. 2009. Genome cyclization as strategy for flavivirus RNA replication. Virus Res 139:230–9.

8. Lindenbach BD, Rice CM. 2003. Molecular biology of flaviviruses. Adv Virus Res 59:23–61.

9. Dong G, Mao Q, Xia W, Xu Y, Wang J, Xu L, Jiang F. 2016. PKM2 and cancer: The function of PKM2 beyond glycolysis (Review). Oncol Lett 11:1980–1986.

10. Dejure FR, Eilers M. 2017. MYC and tumor metabolism: chicken and egg. 36:3409–3420.

11. Liu C, Liu C, Fu R. 2022. Research progress on the role of PKM2 in the immune response. Front Immunol 13:936967.

12. Iqbal MA, Siddiqui FA, Gupta V, Chattopadhyay S, Gopinath P, Kumar B, Manvati S, Chaman N, Bamezai RN. 2013. Insulin enhances metabolic capacities of cancer cells by dual regulation of glycolytic enzyme pyruvate kinase M2. Mol Cancer 12:72.

13. Li Q, Zhang D, Chen X, He L, Li T, Xu X, Li M. 2015. Nuclear PKM2 contributes to gefitinib resistance via upregulation of STAT3 activation in colorectal cancer. Sci Rep 5:16082.

14. Azoitei N, Becher A, Steinestel K, Rouhi A, Diepold K, Genze F, Simmet T, Seufferlein T. 2016. PKM2 promotes tumor angiogenesis by regulating HIF-1α through NF-κB activation. Molecular Cancer 15:3.

15. Liang J, Cao R, Wang X, Zhang Y, Wang P, Gao H, Li C, Yang F, Zeng R, Wei P, Li D, Li W, Yang W. 2017. Mitochondrial PKM2 regulates oxidative stress-induced apoptosis by stabilizing Bcl2. Cell Res 27:329–351.

16. Palsson-McDermott EM, Curtis AM, Goel G, Lauterbach MA, Sheedy FJ, Gleeson LE, van den Bosch MW, Quinn SR, Domingo-Fernandez R, Johnston DG, Jiang JK, Israelsen WJ, Keane J, Thomas C, Clish C, Vander Heiden M, Xavier RJ, O’Neill LA. 2015. Pyruvate kinase M2 regulates Hif-1α activity and IL-1β induction and is a critical determinant of the warburg effect in LPS-activated macrophages. Cell Metab 21:65–80.

17. Shirai T, Nazarewicz RR, Wallis BB, Yanes RE, Watanabe R, Hilhorst M, Tian L, Harrison DG, Giacomini JC, Assimes TL, Goronzy JJ, Weyand CM. 2016. The glycolytic enzyme PKM2 bridges metabolic and inflammatory dysfunction in coronary artery disease. J Exp Med 213:337–54.

18. Andersson U, Wang H, Palmblad K, Aveberger AC, Bloom O, Erlandsson-Harris H, Janson A, Kokkola R, Zhang M, Yang H, Tracey KJ. 2000. High mobility group 1 protein (HMG-1) stimulates proinflammatory cytokine synthesis in human monocytes. J Exp Med 192:565–70.

19. Yang L, Xie M, Yang M, Yu Y, Zhu S, Hou W, Kang R, Lotze MT, Billiar TR, Wang H, Cao L, Tang D. 2014. PKM2 regulates the Warburg effect and promotes HMGB1 release in sepsis. Nature Communications 5:4436.

20. Day AS, Judd T, Lemberg DA, Leach ST. 2012. Fecal M2-PK in children with Crohn’s disease: a preliminary report. Dig Dis Sci 57:2166–70.

21. Chung-Faye G, Hayee B, Maestranzi S, Donaldson N, Forgacs I, Sherwood R. 2007. Fecal M2- pyruvate kinase (M2-PK): a novel marker of intestinal inflammation. Inflamm Bowel Dis 13:1374–8.

22. Tang Q, Ji Q, Xia W, Li L, Bai J, Ni R, Qin Y. 2015. Pyruvate kinase M2 regulates apoptosis of intestinal epithelial cells in Crohn’s disease. Dig Dis Sci 60:393–404.

23. Gupta V, Bamezai RN. 2010. Human pyruvate kinase M2: a multifunctional protein. Protein Sci 19:2031–44.

24. Ren L, Zhang W, Zhang J, Zhang J, Zhang H, Zhu Y, Meng X, Yi Z, Wang R. 2021. Influenza A Virus (H1N1) Infection Induces Glycolysis to Facilitate Viral Replication. Virol Sin 36:1532–1542.

25. McElvaney OJ, McEvoy NL, McElvaney OF, Carroll TP, Murphy MP, Dunlea DM, O NC, Clarke J, O’Connor E, Hogan G, Ryan D, Sulaiman I, Gunaratnam C, Branagan P, O’Brien ME, Morgan RK, Costello RW, Hurley K, Walsh S, de Barra E, McNally C, McConkey S, Boland F, Galvin S, Kiernan F, O’Rourke J, Dwyer R, Power M, Geoghegan P, Larkin C, O’Leary RA, Freeman J, Gaffney A, Marsh B, Curley GF, McElvaney NG. 2020. Characterization of the Inflammatory Response to Severe COVID-19 Illness. Am J Respir Crit Care Med 202:812–821.

26. Zheng D, Jiang Y, Qu C, Yuan H, Hu K, He L, Chen P, Li J, Tu M, Lin LJTAjop. 2020. Pyruvate kinase M2 tetramerization protects against hepatic stellate cell activation and liver fibrosis. 190:2267–2281.

27. Mucaj V, Shay JES, Simon MC. 2012. Effects of hypoxia and HIFs on cancer metabolism. International Journal of Hematology 95:464–470.

28. Bluemlein K, Grüning NM, Feichtinger RG, Lehrach H, Kofler B, Ralser M. 2011. No evidence for a shift in pyruvate kinase PKM1 to PKM2 expression during tumorigenesis. Oncotarget 2:393–400.

29. Cairns RA, Harris IS, Mak TW. 2011. Regulation of cancer cell metabolism. Nat Rev Cancer 11:85–95.

30. Christofk HR, Vander Heiden MG, Harris MH, Ramanathan A, Gerszten RE, Wei R, Fleming MD, Schreiber SL, Cantley LC. 2008. The M2 splice isoform of pyruvate kinase is important for cancer metabolism and tumour growth. Nature 452:230–233.

31. Chen X, Chen S, Yu D. 2020. Protein kinase function of pyruvate kinase M2 and cancer. Cancer Cell International 20:523.

32. Zwerschke W, Mazurek S, Massimi P, Banks L, Eigenbrodt E, Jansen-Dürr P. 1999. Modulation of type M2 pyruvate kinase activity by the human papillomavirus type 16 E7 oncoprotein. Proc Natl Acad Sci U S A 96:1291–6.

33. Mazurek S, Zwerschke W, Jansen-Dürr P, Eigenbrodt E. 2001. Effects of the human papilloma virus HPV-16 E7 oncoprotein on glycolysis and glutaminolysis: role of pyruvate kinase type M2 and the glycolytic-enzyme complex. Biochem J 356:247–56.

34. Wu X, Zhou Y, Zhang K, Liu Q, Guo D. 2008. Isoform-specific interaction of pyruvate kinase with hepatitis C virus NS5B. FEBS Lett 582:2155–60.

35. Zhou D, Jia F, Li Q, Zhang L, Chen Z, Zhao Z, Cui M, Song Y, Chen H, Cao S, Ye J. 2018. Japanese Encephalitis Virus NS1’ Protein Antagonizes Interferon Beta Production. Virol Sin 33:515–523.

36. Mackenzie JM, Jones MK, Young PR. 1996. Immunolocalization of the dengue virus nonstructural glycoprotein NS1 suggests a role in viral RNA replication. Virology 220:232–40.

37. Lindenbach BD, Rice CM. 1997. trans-Complementation of yellow fever virus NS1 reveals a role in early RNA replication. J Virol 71:9608–17.

38. Lindenbach BD, Rice CM. 1999. Genetic interaction of flavivirus nonstructural proteins NS1 and NS4A as a determinant of replicase function. J Virol 73:4611–21.

39. Khromykh AA, Sedlak PL, Westaway EG. 2000. cis- and trans-acting elements in flavivirus RNA replication. J Virol 74:3253–63.

40. Muylaert I, Chambers TJ, Galler R, Rice CM. 1996. Mutagenesis of the N-linked glycosylation sites of the yellow fever virus NS1 protein: effects on virus replication and mouse neurovirulence. Virology 222 1:159–68.

41. Muylaert IR, Galler R, Rice CM. 1997. Genetic analysis of the yellow fever virus NS1 protein: identification of a temperature-sensitive mutation which blocks RNA accumulation. J Virol 71:291–8.

42. Butrapet S, Huang CY, Pierro DJ, Bhamarapravati N, Gubler DJ, Kinney RM. 2000. Attenuation markers of a candidate dengue type 2 vaccine virus, strain 16681 (PDK-53), are defined by mutations in the 5’ noncoding region and nonstructural proteins 1 and 3. J Virol 74:3011–9.

43. Liu X, Cao S, Zhou R, Xu G, Xiao S, Yang Y, Sun M, Li Y, Chen H. 2006. Inhibition of Japanese encephalitis virus NS1 protein expression in cell by small interfering RNAs. Virus Genes 33:69–75.

44. Suzuki R, de Borba L, Duarte dos Santos CN, Mason PW. 2007. Construction of an infectious cDNA clone for a Brazilian prototype strain of dengue virus type 1: characterization of a temperature-sensitive mutation in NS1. Virology 362:374–83.

45. Khromykh AA, Sedlak PL, Guyatt KJ, Hall RA, Westaway EG. 1999. Efficient trans-complementation of the flavivirus kunjin NS5 protein but not of the NS1 protein requires its coexpression with other components of the viral replicase. J Virol 73:10272–80.

